# SEAL: Spatially-resolved Embedding Analysis with Linked Imaging Data

**DOI:** 10.1101/2025.07.19.665696

**Authors:** Simon Warchol, Grace Guo, Johannes Knittel, Dan Freeman, Usha Bhalla, Jeremy L Muhlich, Peter K. Sorger, Hanspeter Pfister

## Abstract

Dimensionality reduction techniques help analysts make sense of complex, high-dimensional spatial datasets, such as multiplexed tissue imaging, satellite imagery, and astronomical observations, by projecting data attributes into a two-dimensional space. However, these techniques typically abstract away crucial spatial, positional, and morphological contexts, complicating interpretation and limiting insights. To address these limitations, we present **SEAL**, an interactive visual analytics system designed to bridge the gap between abstract 2D embeddings and their rich spatial imaging context. **SEAL** introduces a novel hybrid-embedding visualization that preserves image and morphological information while integrating critical high-dimensional feature data. By adapting set visualization methods, **SEAL** allows analysts to identify, visualize, and compare selections—defined manually or algorithmically—in both the embedding and original spatial views, facilitating a deeper understanding of the spatial arrangement and morphological characteristics of entities of interest. To elucidate differences between selected sets of items, **SEAL** employs a scalable surrogate model to calculate feature importance scores, identifying the most influential features governing the position of objects within embeddings. These importance scores are visually summarized across selections, with mathematical set operations enabling detailed comparative analyses. We demonstrate **SEAL**’s effectiveness and versatility through three case studies: colorectal cancer tissue analysis with a pharmacologist, melanoma investigation with a cell biologist, and exploration of sky survey data with an astronomer. These studies underscore the importance of integrating image context into embedding spaces when interpreting complex imaging datasets. Implemented as a standalone tool while also integrating seamlessly with computational notebooks, **SEAL** provides an interactive platform for spatially informed exploration of high-dimensional datasets, significantly enhancing interpretability and insight generation.

## 1 Introduction

High-dimensional spatial data—e.g., multiplexed tissue, satellite, and astronomical imagery—encode multi-channel information and spatial relationships. For instance, in multiplexed tissue imaging, each pixel in a high-resolution microscopy image may represent a spot within a cell along with dozens of associated biomarker intensities, while in multispectral astronomical imagery, each pixel may contain multiple spectral bands capturing subtle differences in surface or atmospheric properties [1, 4, 16, 42]. Large-scale, high-dimensional datasets offer new insights across domains—from tumor–immune interactions in oncology [61] to galaxy evolution in astrophysics [11, 14]—but remain challenging to analyze due to their complexity.

To make these data more interpretable, dimensionality reduction (DR) methods such as PCA [63], t-SNE [48], and UMAP [54] are essential. By projecting high-dimensional features into two dimensions, DR methods enable researchers to visualize complex structures and identify clusters, outliers, and other meaningful patterns [6, 32].In many domains, scatterplots of these 2D embeddings have become a standard tool to guide exploratory analyses, enabling experts to identify cell types in tissue imaging or classify celestial objects in astronomy [11, 61]. To investigate how underlying features are represented within embeddings, points in these plots are generally colored by their feature values (Fig 2). However, while such plots reveal valuable high-level patterns, they abstract away the critical spatial details in the original images. Furthermore, while some prior studies [19, 30, 55, 91] explore possibilities for using positional data to enrich low-dimensional embeddings, spatial characteristics are more than just positional and can include shape, area, color, channel intensity, and neighboring regions. For instance, embeddings of single-cell data extracted from larger tissue images (e.g., Fig 3) neglect the broader architectural context—such as where in the tissue a cell resides, its morphology, and the proteins it expresses. Thus, bridging the gap between these abstract 2D embeddings and the rich spatial context of the original data remains a major open challenge.

**Figure 1.**
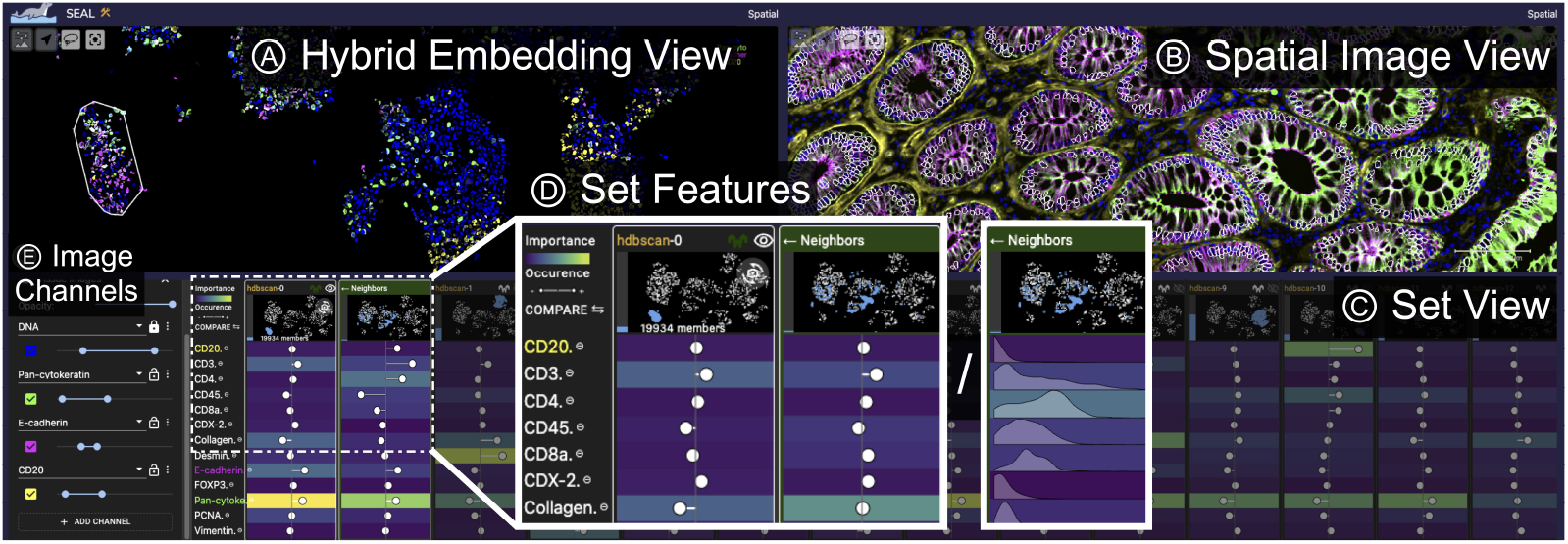
**SEAL** visualizing a colorectal cancer specimen and its corresponding single-cell embedding. A) The Hybrid Embedding View integrates multi-channel imaging data and entity morphology into a 2D projection. B) The Spatial Image View preserves positional context and reveals tissue organization. C) The Set View highlights identified sets along with their distinguishing high-dimensional features, D) visualizing each set’s embedding position, feature importance, occurrence, nearest neighbors, and, optionally, data distribution. E) Image channels can be pseudocolored and visualized simultaneously. Here, a cluster in the embedding corresponds to epithelial crypts in the tissue image.

**Figure 2.**
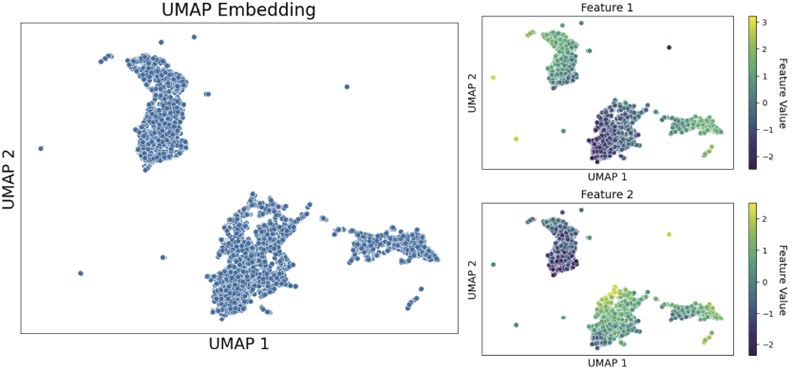
Embedding and Feature Visualization: Scatterplots are commonly used to visualize embeddings (left), and to explore how the raw values of individual features (right, encoded with color) vary across clusters. Here, two arbitrary features are shown for illustration.

**Figure 3.**
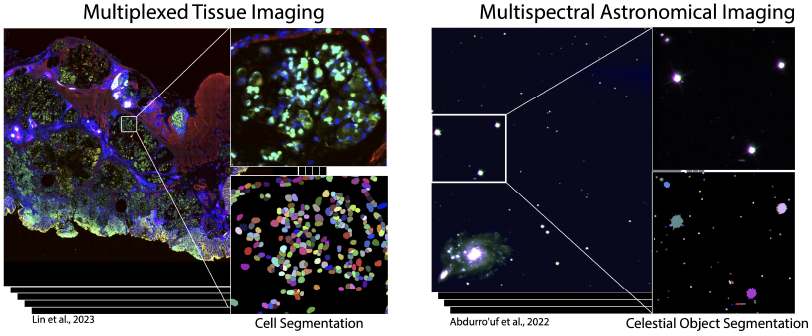
Spatial Imaging Data: **SEAL** supports expert analysis of multi-channel spatial images, including multiplexed tissue samples and multi-spectral astronomical data that contain segmented entities (e.g., cells, stars, galaxies).

To address this, we present **SEAL**, a novel visualization approach that links multichannel imaging data and 2D embeddings. We make the following contributions: **(1)** a novel multi-resolution Hybrid Embedding visualization that places segmented entities—such as cells or celestial objects—into the embedding space, retaining both their morphology and image channels. Each entity is represented by a cutout from the original image, and a semantic zoom mechanism reveals additional entities at higher resolutions, avoiding occlusion while preserving spatial detail. **(2)** To better connect the Hybrid Embedding with the original high-dimensional data, we employ a scalable surrogate model to compute per-entity feature importance scores. These scores help explain the position of each point in the embedding, enabling users to explore and interpret the underlying features that shape its structure. **(3)** Users can compare clusters or ROIs within the embedding or original image through a set-comparison approach, allowing them to identify key features, such as biomarker compositions, that distinguish selections of cells from one another. **(4)** We incorporate these functions into a combined visual analytics tool, **SEAL**, which was developed to support the above tasks and can be used as a standalone application or embedded in computational notebooks. **(5)** We demonstrate the utility of **SEAL** through three case studies: two with cancer researchers working to understand the composition of the tumor microenvironment and one with an astronomer investigating relationships between celestial bodies in a portion of the Sloan Digital Sky Survey [1].

## 2 Related Work

A substantial body of research has addressed the visualization and interpretation of high-dimensional spatial data, particularly where spatial structure, multivariate features, and group comparisons are key.

### 2.1 Dimensionality Reduction

Some of the most widely used DR algorithms include PCA [63], t-SNE [48], and UMAP [54]. These methods involve trade-offs, particularly in preserving local versus global structure; linear methods like PCA capture global structure but miss local detail, while nonlinear methods like t-SNE and UMAP do the reverse [33]. Such trade-offs influence how users interpret local neighborhoods in relation to broader cluster structure. Various algorithms mitigate these trade-offs in distinct ways: Isomap [79] preserves global geometry via geodesic distances, TopoAE [58] enforces topology through a persistent-homology loss, UMATO [26] aligns local and global structure using hub anchors. Hierarchical methods like HSNE [64] and HUMAP [51] preserve neighborhood structure across scales via multi-resolution embeddings. Recent work has also incorporated spatial or texture features directly into DR, improving the embedding of imaging data [83]. New metrics have been proposed to evaluate how well DR methods preserve clusters from the original high-dimensional space [24, 25, 27, 60, 81]. **SEAL** is algorithm-agnostic, enabling users to choose a DR method best suited to their goals or investigate embeddings from prior studies.

### 2.2 Embedding Visualization and Analysis

Scatterplots are a popular and effective tool for exploring 2D embeddings, with a wide range of methods developed to further improve their clarity and interpretability.

#### Addressing Overplotting

Overplotting is a common challenge when visualizing large datasets with scatterplots; approaches such as Hagrid [17], GIST [36], Rave et al. [68], and Quadri et al. [67] adjust point position, opacity, or sampling to better preserve density and cluster structure. Splatterplots [53] reduce visual clutter by abstracting dense regions into smooth, contour-like shapes, maintaining key patterns, and highlighting outliers. In contrast, Manz et al. [50] transform labeled embeddings to accentuate distinctions between classes, thereby making inter-class differences more perceptible.

#### Adding Annotations

Annotations enrich embedding visualizations by adding semantic context and clarifying data structure. Wang et al. [86] use contours and hierarchical labels to highlight concept density and relationships, while Li et al. [41] group points into bounded, labeled regions based on class. Other approaches [48, 79] display images at embedding locations to visually validate structure, an approach closely related to our Hybrid Embedding. In **SEAL**, we build on both approaches, adding density contours and supplementary set labels that include feature importance information. At the same time, we preserve the original embedding’s structure and introduce an occlusion-removing semantic-zoom approach that incorporates image context and entity morphology, thereby providing rich, context-aware cues that enhance the perceptibility of inter-class relationships.

#### Interactive Approaches

Despite their popularity, 2D embeddings of high-dimensional data are hard to interpret, especially when plots lack clear cluster separation, leading to subjective analysis [28]. Interactive tools help users explore and interpret embeddings more effectively. Interactive hierarchical DR techniques, such as HSNE [64] and HUMAP [51], employ *Overview-First, Details-On-Demand* workflows to support exploration while preserving spatial continuity. Splatterplots [53] progressively reveal abstracted data details through zoom-based interactions, managing visual complexity. Vieth et al. [82] link hierarchical embeddings with image navigation for deeper exploration. DimBridge, meanwhile, links visual patterns in the scatterplot to the underlying features [57], while Corbugy et al. chose to link feature importance plots to scatterplots to help users identify how elements differ within and between clusters [15]. Sohns et al. [76] visually group data points using geometric annotations, highlighting local structure and enabling interactive exploration. Jeon et al. [23] introduce distortion-aware brushing to dynamically correct misleading relationships in embeddings. Embeddings that represent images or volumes benefit significantly from interpolations between points, further enhancing interpretability [8, 44].

We do not aim to recover information lost in projecting high-dimensional embeddings to two dimensions. Instead, **SEAL** enriches embedding clusters by linking them to multi-channel spatial representations and enabling bidirectional exploration. Prior techniques approach this by coordinating multiple views—using color to reveal cluster compositions [50], attaching neighborhood metrics [21, 40], or revealing local structure via small multiples [7]. Other tools place images directly in the embedding [12, 47, 52, 65]. **SEAL** builds on these advances by unifying spatial imaging data with high-dimensional feature analysis, fusing morphology and position with feature-importance cues that help users explain both the embedding and its spatial entities.

### 2.3 Visualization of Multivariate Spatial Data and Clusters

Effective visualization of multivariate spatial data requires linking spatial context to high-dimensional attributes, thus revealing patterns, relationships, and clusters across dimensions.

#### Graph Overlays

Integrating spatial and multidimensional data has been widely studied in geographic information systems (GIS). Common approaches overlay glyphs and/or concave hulls on top of spatial maps to define, inspect, and compare geographic clusters. For example, Park et al. place multivariate glyphs at varying spatial granularities [62], while ConcaveCubes [39] and TopoGroups [90] introduce methods for clustering and visualizing concave hulls on maps.

#### Linking Views

Linking scatterplots to multiplexed tissue images and segmentation masks is common in single-cell analysis [35, 73, 77, 87]. More broadly, coordinated multi-view systems combine maps with complementary visualizations for added context. For example, CriPAV links maps, embeddings, and street views for crime analysis [19], while SNoMAN connects maps and graphs for spatial social network exploration [30]. Urban Mosaic supports similarity search across street views and maps, without explicit embedding visualization [55], and Zhang et al. [91] examine how spatial variation affects clustering outcomes. Previous work typically emphasized positional information; however, spatial characteristics also encompass shape, size, local composition, and, for images, pixel-level attributes such as color and intensity.

### 2.4 Set Visualization

Most visualization approaches for 2D embeddings treat *clusters* as the key unit of analysis. They aim to visually distinguish clusters in the scatterplot [17, 23, 28, 36, 50, 67, 68], describe the constituent data items in each cluster [15, 41, 50, 57, 86], and compare clusters to identify similarities and differences [15, 50]. We extend these approaches by treating clusters and user-defined selections as *sets*. Set visualizations provide flexible ways to depict and compare subgroups in data. Techniques like Bubble Sets [13] and inverse distance-based methods [84] use contours to highlight groups, while Kelp Diagrams [18] and LineSets [2] connect group members directly. Some methods, such as those by Simonetto et al. [74], jointly optimize layout and set boundaries. Grid-based techniques like OnSet [70] and GridSet [10] use discrete units for representing set operations. Others, such as UpSet [38], employ chart-inspired and radial or flow-based layouts [3, 34] to convey overlapping memberships and relationships.

While set visualizations effectively show group membership, they often overlook underlying feature distributions and spatial attributes such as shape, area, color, and intensity [2, 13, 18, 84], limiting their value for multivariate spatial analysis. To address this, **SEAL** introduces a Hybrid Embedding that integrates multi-channel image cutouts into 2D plots, enabling pattern exploration with pixel-level detail. Building on prior methods, **SEAL** treats selections as sets—derived from annotations or clustering—and uses feature importance scores to characterize and compare these sets via *union, intersection*, and *difference*.

## 3 Data Description

**SEAL** is designed to support a wide range of scientific imaging datasets that are both high-dimensional in their feature representations and rich in spatial and morphological structure (Fig 3). Examples include multiplexed tissue images with dozens of molecular markers per cell [16, 42], multispectral or hyperspectral satellite imagery [4], and wide-field astronomical surveys capturing millions of celestial objects across multiple wavelengths [1]. Experts typically explore such data by zooming, panning, and pseudocoloring to highlight localized patterns, but this makes it difficult to discern global trends across large datasets.

**SEAL** addresses this challenge by operating on datasets with four key characteristics: (1) a multi-channel image in which each pixel contains multiple measurements, (2) segmentation masks that identify individual entities, (3) per-object features such as mean intensities or morphological descriptors, and (4) a two-dimensional embedding generated via dimensionality reduction methods such as UMAP, t-SNE, or PCA. This approach assumes that discrete entities—such as cells, stars, or geographic regions—can be segmented from the image. While dimensionality reduction helps reveal structure in high-dimensional data, traditional scatterplots discard spatial and morphological context, making it difficult to interpret embeddings in relation to the original image. **SEAL** overcomes this by linking each embedded entity back to its spatial footprint and feature values, preserving context critical for interpretation. For instance, in single-cell pathology, embeddings may reveal cellular clusters but obscure their spatial organization within a tumor [61,85]. As shown in Fig 3, these requirements are satisfied in diverse domains including cancer tissue profiling [16, 42, 61], astronomical surveys [66], and satellite imaging [20, 78], making **SEAL** broadly applicable across disciplines.

## 4 Task Requirements

To understand how domain experts analyze spatially resolved datasets, we conducted feedback sessions using early prototypes of **SEAL**. Over the course of a year, we held meetings with approximately 20 researchers working with spatial data—including those in oncology, cell biology, and astrophysics. Our discussions revealed that, despite differences in subject matter, experts in both domains follow similar workflows: they iteratively generate 2D embedding scatterplots (often within Jupyter notebooks) for hypothesis testing, then map variables to color or opacity to assess their impact on the embedding’s structure, as demonstrated in Fig 2. However, because the DR strips away critical spatial and imaging context, the choice of which variable to encode in the scatterplot is often guided only by prior knowledge or best guesses. This makes it challenging to correlate observed patterns with underlying data attributes. Our collaborators emphasized the importance of maintaining spatial context, noting that clusters in embedding spaces frequently mirror complex spatial structures in the original data and that visually distinct regions can reveal significant relationships. This shared approach underscores not only the similarities in workflows across domains but also the pivotal role of embeddings in driving meaningful insights.

Our discussions highlighted the researchers’ strong interest in achieving a spatially informed interpretation of embedding spaces. Analysts recognize that clusters found in the embedding space often map to complex spatial structures in the original dataset, just as visually interesting spatial regions can exhibit shared attributes within the embedding space. Motivated by feedback from our initial meetings and from our prior work supporting their analysis workflows [29, 35, 87, 88], as well as the broader needs of the highdimensional spatial data domain [9, 37], we identified four core analysis tasks to guide the design of **SEAL**. We aim to provide means for experts to **situate, select, characterize**, and **compare** patterns across both spaces. Note that we use the term “entity” to refer to a segmented object in the dataset (e.g., a cell or galaxy), and “set” to refer to any selection of such entities—whether defined algorithmically (e.g., a cluster) or manually (e.g., a region of interest) in the 2D embedding or the original spatial image.

### T1: Situate data in both the 2D embedding and spatial image

The 2D embedding highlights patterns in high-dimensional attribute space, while the spatial image shows patterns based on physical location. Users should be able to quickly understand how each data item is positioned in both views; visual patterns identified in one view should highlight the pattern composed of the same data items in the other view. Indeed, this involves not only comparing clusters in the embedding, but also relating a given cluster to a spatial region of interest (ROI), as well as comparing patterns across multiple ROIs. For example, an expert could identify a cluster in the embedding that represents a certain immune cell type. The corresponding spatial position of cells in that cluster might then show how these immune cells are arranged to surround tumor masses.

### T2: Select sets of data items based on different criteria

Since the embedding and the spatial image reveal different data patterns, users should be able to make selections from the two views based on different criteria. The embedding, for example, lends itself well to algorithmic clustering approaches, such as k-means, that can automatically identify similar data items. In contrast, the spatial image may reveal mesoscale features or gestalt patterns that are interesting to human domain experts. Similarly, neighbors in the embedding space indicate data items with similar multivariate attributes, while neighbors in the spatial image indicate entities that physically interact or are otherwise morphologically related. To account for these different ways of identifying sets of interest, the tool should thus support different methods of selecting items from the data.

### T3: Characterize sets in terms of their distinguishing features and attributes

This provides a concise description of each selected set, and helps users understand the distinguishing features of its constituent data items. For example, given a selected cluster of cells, a cell biologist should be able to identify key features—such as biomarkers—that are most correlated with membership, revealing key biological properties.

### T4: Compare multiple sets of items

Comparing sets is crucial for helping users build a nuanced understanding of how sets are similar and different across their spatial, morphological, and highdimensional characteristics. For example, a cell biologist may select two different groups of immune cell types from the embedding space. The intersection of these two sets would reveal features or biomarkers that are common across both cell types, while the difference of these two sets would reveal distinguishing features for each. To support more complex comparisons, users should be able to quickly create new subsets from prior comparisons (e.g., the intersection of two sets becomes a new subset of items).

## 5 SEAL Visualization Approach

To facilitate the exploration and analysis of embeddings and their corresponding spatial images, we introduce **SEAL** (Fig 1), a novel visual analytics system designed to integrate spatial, morphological, and feature-based context into embedding visualizations. **SEAL** contains three main views: 1) a **Hybrid Embedding** view, 2) the original **Spatial Image** view, and 3) the **Set View**, which displays the feature importance and spatial distribution of all selected sets. The views are linked and designed to address the task requirements in Section 4.

### 5.1 Adding Context to Embeddings

Standard scatterplot embeddings offer an efficient way to visualize high-dimensional patterns, but they often obscure the spatial and morphological characteristics critical for interpretation. As identified in our task analysis (Sec 4), experts frequently need to **situate (T1)** data to understand how patterns in the embedding correspond to spatial structures in the original image and vice versa.

To address this need, we developed the **Hybrid Embedding** view, which augments traditional embeddings with imaging context. Each data point in this view corresponds to a single segmented entity; we display the corresponding multi-channel image patch, cut out from the original image using its segmentation mask, directly at its location in the embedding space. These image cutouts function as glyphs, providing a direct visual representation of entity shape, structure, and expression across channels. Generating this view—and the **SEAL** approach more broadly—assumes that the input image contains a large number of small, similarly sized entities (e.g., cells, stars) that can be reliably segmented and extracted, forming the basis for spatially contextualized embedding visualizations.

We begin with the original **Spatial Image** view (Fig 4a), from which we cut out the segmented entities using CellCutter^1^, a domain-agnostic tool for separating segmented entities, retaining everything contained within the segmentation mask for each image channel (Fig 4b).

**Figure 4.**
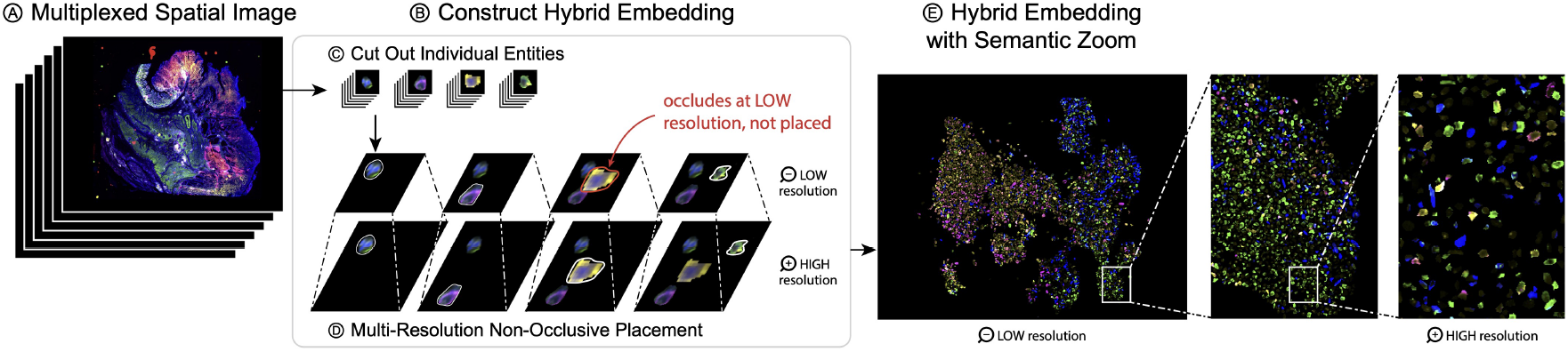
The Hybrid Embedding Image integrates spatial morphology and multiplexed feature data directly into the embedding space. (A) From the original Spatial Image, we (B) construct the hybrid embedding by (C) segmenting and cutting out entities, preserving all pixel information within each segmentation mask. (D) For each segmented entity, we place them in the embedding space at different resolutions. If an item occludes another, it will be omitted from that resolution. (E) This creates the hybrid embedding space where zooming in reveals more entities.

This yields an *NxCxBxB* array, where *N* is the number of entities in an image, *C* is the number of channels, and *BxB* is the size of the largest bounding box for an object in the image. We then construct a new image with the same size as the original and scale the projection accordingly. This new image, like the original, is organized as a multi-resolution pyramid. At each resolution in the image pyramid (Fig 4c), we randomize the order of the objects and place all possible entities in the new image such that no two entities overlap at the given resolution. Entities that cannot be placed without overlap are excluded at a given resolution, effectively occluded from the view, but are progressively revealed as users zoom in and additional space becomes available for non-overlapping placement. The final **Hybrid Embedding** image (Fig 4d) thus supports a semantic zooming effect (Fig 4e), where, as a user zooms in, more and more objects are loaded to provide further detail on the composition and entity morphology within that region. Moreover, as each entity in the embedding retains its multi-channel imaging data, we are able to preserve high-dimensional feature fidelity and entity morphology within a spatially meaningful layout. Experts may thus explore the **Hybrid Embedding** as they would the original multi-channel image. While zooming into the **Spatial Image** reveals the intricate spatial arrangement of entities, zooming into the **Hybrid Embedding** progressively renders additional entities—preserving both their morphology and high-dimensional feature representation—allowing users to examine the composition of the embedding in greater detail. Moreover, because both views display multi-channel image data, users can explore individual entities or regions through pseudocoloring, providing full spatial context for the expression of specific features (see Sec 5.2). Upon selecting a set or individual entity, **SEAL** automatically highlights the most relevant image channels based on feature importance (see Sec 5.5), guiding users toward the channels most likely to reveal informative patterns. Thus, as detailed in subsequent sections, this enables targeted visual exploration by aligning high-dimensional feature relevance with interpretable spatial cues in both the **Spatial Image** and **Hybrid Embedding** views.

### 5.2 Linking the Spatial Image and Hybrid Embedding

Linked analysis of the **Spatial Image** and **Hybrid Embedding** views allows for a more nuanced understanding of the data by situating **(T1)** entities in both spaces. Both views display multi-channel images, where users can select and combine channels. Each channel is pseudo-colored, where intensity values are mapped to a specific color by interpolating between black and the chosen color, enhancing visual contrast and feature distinction. Channel selection and color mapping are synchronized across both the **Hybrid Embedding** and **Spatial Image**, enabling users to navigate seamlessly between views while inspecting morphology and feature expression. Users may also toggle the background color from black to white to accentuate dark entities in either view.

A limitation of our **Hybrid Embedding** approach is that, by deliberately eliminating occlusion, we no longer encode the density of the embedding space. This limitation is shared with scatterplots themselves, and numerous approaches exist to balance accurate data representation, entity preservation, overplotting, and occlusion [72]. However, in our implementation, we found that methods that distort an embedding’s shape scale poorly for datasets with millions of points, especially in dense regions representing large groups of homogeneous entities (e.g., tumor cells). Furthermore, expert feedback suggested it is more effective to display a single exemplar at a given resolution rather than many overlapping entities. For instance, while an expert can identify a specific star type when viewed in isolation, displaying many such galaxies together obscures this finding. As such, we augment the **Hybrid Embedding** in **SEAL** with a minimally invasive contour plot of the embedding density that the user can toggle on or off (Fig 5). To further address the possibility that a given entity of interest may not be visible in the current view, the original scatterplot may be overlaid on the **Hybrid Embedding**, with selected entities encoded using color.

**Figure 5.**
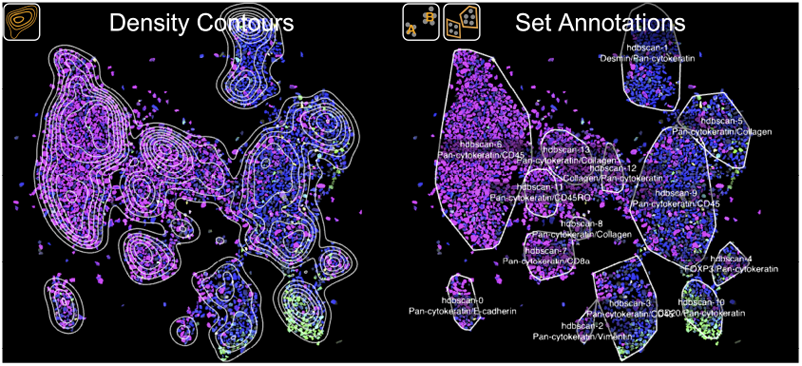
Augmenting the Hybrid Embedding View. Contour lines encode the density of the Hybrid Embedding. Set annotations outline a set with a convex hull and list the most important features to that set.

### 5.3 Selecting and Visualizing Sets

**SEAL** supports multiple methods for defining and **selecting (T2)** sets of data items. Clusters and regions of interest (ROIs) may be defined directly by manually lassoing areas in either the **Hybrid Embedding** view or **Spatial Image** view. This approach permits the identification of clusters in the embedding space as well as the delineation of notable spatial entities. In addition, pre-defined sets—such as those produced by clustering algorithms or derived from prior analyses—may be imported. When operating SEAL within a Jupyter notebook, clustering can be iteratively refined by adjusting algorithms and hyperparameters in real time, all within the same environment. Moreover, since sets may have overlapping membership, SEAL facilitates the comparison of hierarchies or clustering algorithms and the dynamic dissection of existing sets using the lasso tool.

The entities that compose a given set are then highlighted in both the **Hybrid Embedding** view and **Spatial Image**. To minimize occlusion and maximize visibility, we highlight set members in both views without using color, as color encodes feature expression in the multi-channel image. Instead, we employ the techniques of *outlining* and *spotlighting* (Fig 6):

**Figure 6.**
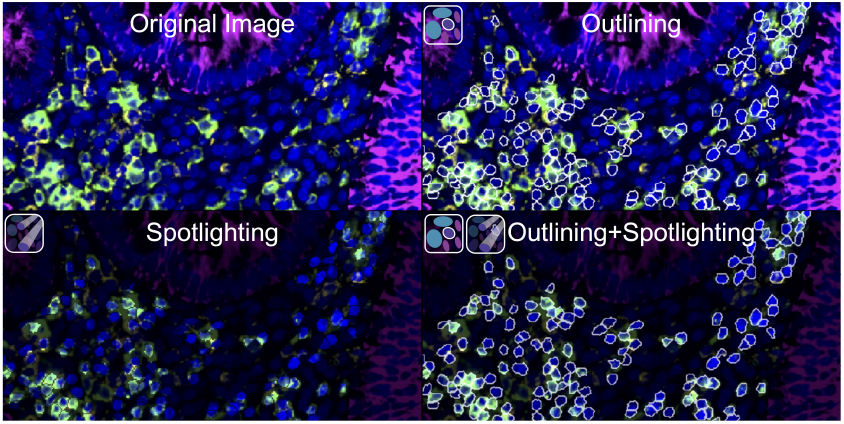
Visualizing sets in multi-channel images: Outlining, Spotlighting, and their combination, enhance entity visibility.

- **Outlining:** The segmentation mask is used to draw a white border around each selected entity. These borders are placed outside the entity to prevent occluding its morphological features.
- **Spotlighting:** The luminance outside each segmented entity is reduced while the original luminance within the entity is maintained. Since the original image comprises both segmented entities and a non-segmented background that may provide additional context, the dimming can be applied to the unselected entities, the background, or both, further highlighting the selected entities.

These two techniques serve distinct yet complementary roles. While both preserve the multi-channel morphological details of each selected entity, spotlighting immediately draws attention to those entities at a global scale, highlighting regions that users may inspect at higher resolution. However, as discussed in Sec 4, the surrounding area and neighboring entities offer critical contextual cues. Thus, outlining selections ensures that this external context remains visible and unaltered.

In addition to highlighting the entities that compose a set, we add annotations to both views to indicate the set membership; when the entities in a set are above a certain density, we optionally draw a convex hull around them (Fig 5). Clusters can also be labeled with their names and most important features (see Sec 5.5), with annotations positioned at their centroid (Fig 5). Consistent with the selection modes, these annotations are carefully designed to minimize occlusion and visual clutter while clearly conveying both the spatial distribution of sets and the distinguishing features of their constituent entities. Moreover, when these annotations correspond to specific image channels, clicking on a set visualizes and pseudocolors these corresponding channels in the **Hybrid Embedding** and **Spatial Image** views, enabling users to verify the annotations and inspect the underlying entities in greater detail.

### 5.4 Exploring Local Context with Set Neighbors

Our prior collaborations with domain experts have emphasized the importance of looking beyond the spatial locations of entities. Indeed, investigating the spatial neighborhoods is crucial to achieving a comprehensive understanding of spatial imaging data [87]. To address this, we introduce a feature in **SEAL** that identifies neighboring entities of a selected set, enabling users to **select** these neighbors **(T2)** as a new set. By default, this uses a k-nearest-neighbors (k-NN) query, with optional *ε*-neighborhood selection for spatial views. While k-NN ensures consistent result sizes, *ε*-neighborhoods offer more intuitive control by selecting all entities within a user-defined radius. This capability is valuable for exploring spatial relationships, identifying peripheral structures, or uncovering subtle patterns in the local context of a set.

### 5.5 Explaining Sets with Feature Importance

While the **Hybrid Embedding** and **Spatial Image** convey high-dimensional patterns and spatial relationships between data items, users may also want to understand the precise attributes that distinguish a selected set and how it fits into the broader embedding structure. To support this, we **characterize** each set and its constituent entities in terms of their distinguishing features **(T3)**.

To achieve this, we employ SHapley Additive exPlanations (SHAP) values [46], a theoretically grounded method based on cooperative game theory for interpreting the output of predictive models. SHAP values provide an intuitive, unified measure of feature importance, quantifying how each feature contributes individually to the predictions [56], with contributions summing to the model’s output. Unlike traditional methods such as gain or permutation importance, SHAP ensures consistency, guaranteeing that increasing a feature’s true impact does not decrease its assigned importance [45]. This property is critical for reliable interpretation, particularly when used with complex models like XGBoost, and has led to its wide adoption across domains [22].

Since UMAP and t-SNE do not expose how original features drive the 2D layout, we train a global surrogate model mapping the high-dimensional inputs to their embedding coordinates [56]. Following Molnar’s “Global Surrogate” recipe, we assemble (features, embedding) pairs and grid-search XGBoost hyperparameters, limiting depth to six, to maximize surrogate *R*^2^ on held-out data as our fidelity metric [56]. We then confirm that the same threshold (*≥*0.95) holds for both *R*^2^ and the trustworthiness score, ensuring strong global and local agreement in our case studies [80]. Note that these thresholds are specific to our test and case study embeddings, reflecting both embedding quality and their predictability from the original features. Researchers replicating this workflow should combine these quantitative metrics with manual inspection of the reconstructed embedding and may adopt any fidelity or importance measures they deem appropriate. Finally, we extract per-point SHAP values via Tree SHAP, which **SEAL** ingests in Parquet format. While our proposed global surrogate produced meaningful attributions and novel insights (see Sec. 7.1), we acknowledge the value of alternative methods. Users may compute feature importance by any preferred approach, e.g., gradient based attributions on a parametric UMAP embedding [71], model agnostic surrogates like LIME [69], or custom techniques, and supply those values to **SEAL** in the expected format.

We visualize the mean per-feature SHAP values for each set in the **Set View** as a heatmap using the viridis color map [75] (Fig. 1d), essentially representing the average impact of each feature on the set’s embedding positions. Since high importance can reflect either presence or absence, we overlay a lollipop plot indicating relative feature incidence in the set—lollipops extending right denote features more prevalent in the set than in the full dataset, and vice versa. Users may switch encodings (e.g., a diverging color map for incidence and lollipops for importance) and sort sets by feature. As an alternative to the lollipop plots, users can display a density plot of SHAP or incidence values to reveal internal variability, which is useful when selections contain distinct subpopulations that may be obscured by averaging (see Fig. 1d). If a feature corresponds to a **Spatial Image** channel, selecting a set updates the view to pseudocolor the most distinguishing channels (unless locked). Above each set, toggleable thumbnails (between **Hybrid Embedding** and **Spatial Image**) provide a quick view of spatial distribution and clustering (Fig. 1d), with a bar next to each thumbnail showing set size to convey density information absent from the embedding.

### 5.6 Comparing Sets

To support users in performing set **comparisons (T4)**, the **Set View** presents an integrated heatmap visualization that consolidates per-set feature importance and relative feature incidence (Sec 5.5), facilitating intuitive visual assessment of feature variation across sets. This view assists users in investigating similarities and differences among selected sets—whether chosen within the **Hybrid Embedding**, within the **Spatial Image**, or simultaneously from both views. These comparisons are crucial for understanding how sets differ in the high-dimensional feature space.

Beyond this high-level visual comparison, we also allow users to conduct more complex analysis of feature importance between sets in a pairwise manner. Many prior works on set visualization include some mathematical set operations, most commonly by displaying set intersections and unions [10, 38, 70]. We adapt this feature in **SEAL** by extending our support to all available mathematical set operations (*union, intersection, difference, symmetric difference*, and *complement*). When a user selects two sets, **SEAL** will display each of the mathematical set operations as a new set in the comparison view, color-coded based on the operation (Fig 7).

**Figure 7.**
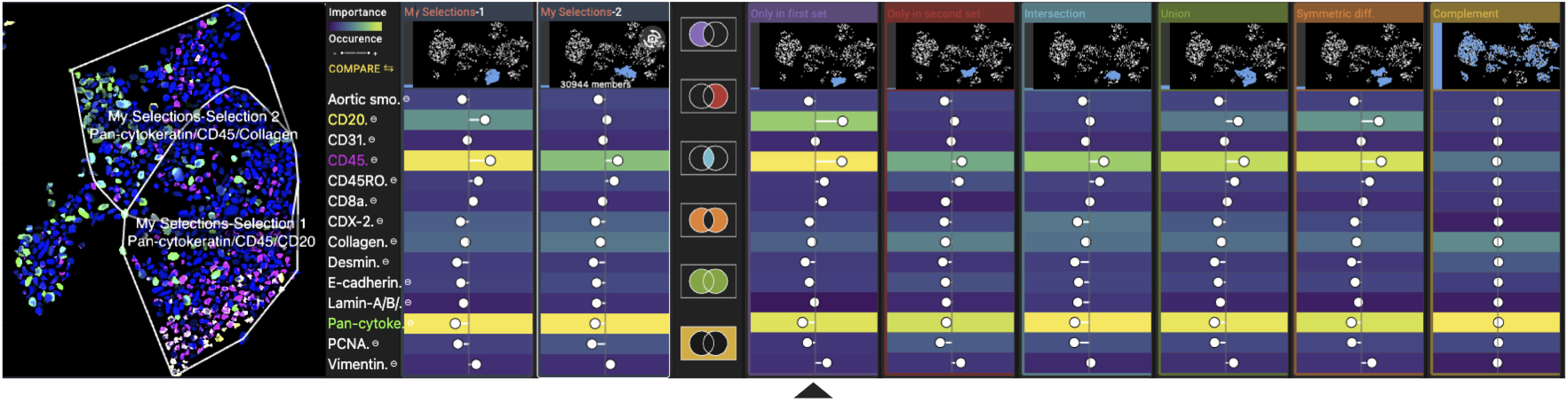
Set Comparison. Two overlapping ROIs from a larger cluster identified in Case Study 1 in the Set Comparison View. A discrete population of immune cells is revealed by the “Only in first set” operation, as indicated by the arrow. This set is distinguished by the presence and importance of CD45 (an immune biomarker) and CD20 (a B cell biomarker).

We exclude any set operations that yield no meaningful information. For example, we omit intersections between sets with no shared elements, such as *A* ∩ *B* when *A* and *B* have no overlap. Similarly, we do not display set differences like *A \B* or *B \A*, as these simply reproduce each set in isolation within this context. Instead, we focus on the remaining operations that provide more interpretive value. The *union A* ∪ *B*, in the **Set View**, emphasizes the features of greatest shared importance between both sets. Conversely, the *complement of the union* (*A* ∪ *B*)′ reveals where entities not present in either set reside, along with their associated feature importance.

When sets share members, their *intersection A* ∩ *B* captures the shared entities and presents them as a new set in the **Set View**. This allows users to investigate commonalities in feature importance and incidence, shedding light on shared functional or structural properties. For example, selecting two spatially distinct regions of tumor cells may reveal an intersection with features characteristic of tumor biology in general—such as elevated proliferation markers or immune evasion signatures—thus offering insights into shared traits across regions.

Conversely, operations that highlight differences between sets reveal features unique to each, helping to explain functional or phenotypic divergence. Continuing the tumor comparison example, differing features between two sets may point to variations in immune infiltration, tumor microenvironment, or therapeutic response. Such differences are critical for understanding heterogeneity within or across regions. This supports both exploratory and confirmatory workflows: users might compare a spatial region of interest (ROI) against a cluster from the embedding to investigate how spatial localization aligns with high-dimensional feature space, or analyze how a subset differs from its parent set in terms of expression patterns, morphology, or cell state.

## 6 Implementation

**SEAL** is implemented both as a standalone web application and as a Jupyter widget. The visualization components are built with Vittessce [31], D3, and Deck.gl. We support multichannel images stored either in the OME-TIFF or OME-ZARR [59] format, which are ideally stored in a cloud bucket. The **SEAL** backend is implemented in Python using FastAPI. We use anywidget [49] to package the Jupyter widget, which allows us to use the same codebase and provide a consistent user experience across the web application and the Jupyter notebook. This allows users to investigate different clustering and DR algorithms interactively within the same notebook and immediately link their outputs back to the original spatial imaging context. The system and example notebooks are publicly accessible at https://sealvis.com. The underlying source code will be made open access upon publication.

## 7 Case Studies

We demonstrate the effectiveness of **SEAL** through three case studies in oncology and astronomy with domain experts.

### 7.1 Colorectal Cancer Analysis with a Pharmacologist

For the first case study, we met with an immuno-oncology researcher with over a decade of experience investigating tumor invasion and immune evasion, now specializing in high-dimensional immuno-profiling to uncover how solid tumors manipulate the immune system. They bring a systems-level perspective to single-cell data, distinguishing true immune signatures from artifacts and informing immunotherapy strategies. The study session lasted an hour, during which the researcher examined a well-curated colorectal cancer specimen (part of a larger cancer atlas [43]) containing 22 protein biomarkers. Cells were segmented [89] and subjected to rigorous quality control via CyLinter [5]. The mean channel intensity values for each segmented cell were then embedded using UMAP, revealing well-separated clusters corresponding to annotated cell types. The dataset is meticulously curated and is thus an ideal test bed for validating **SEAL**.

The researcher first scanned the **Hybrid Embedding**, using the contour visualization to check for hidden high-density regions (Fig 5). Finding none, they switched to the **Set View**, where they observed that in nearly every set, the biomarker pan-cytokeratin showed high feature importance. This was expected, as pan-cytokeratin distinguishes tumor cells from normal cells; thus, its presence in the embedding generally indicates a tumor cell or a tumor-adjacent function. From the heat map alone, they could ascribe several cell types to specific sets. They identified a large, well-differentiated cluster with high feature importance and strong expression of pan-cytokeratin and PCNA (a proliferation marker), combined with negative expression of collagen (an epithelial marker). They concluded this was a tumor-cell cluster.

Moving on, the expert toggled the cluster annotations in the **Hybrid Embedding** view and noticed that the cluster (“hdbscan-3”) containing the typical B-cell markers (Fig 5) was much larger than anticipated. Upon investigating the embedding, it appeared to contain multiple subclusters. They then selected the upper and bottom portions of this cluster using the lasso tool and used the comparison view to investigate this area of heterogeneity (Fig 7). In doing so, they found that the bottom portion of this cluster contained a discrete population of B-cells, whereas the upper portion contained a portion of stromal and T cells.

The researcher then examined a nearby CD163-positive cluster, a biomarker associated with macrophages. Zooming in, the cells showed the expected macrophage morphology: they were larger and had less rounded, more uneven shapes when compared to white blood cells, such as T and B cells. This agreement between marker expression and morphology, visible directly within the **Hybrid Embedding**, reinforced the annotation and demonstrated the value of integrating molecular and spatial information.

The expert was then intrigued by a set forming a well-differentiated cluster that had high feature importance and relative expression of pan-cytokeratin, CD3 (a T-cell marker), and E-cadherin (a marker for normal epithelium)—indicating seemingly conflicting cell types or functions. Upon examining these cells in the **Spatial Image**, they were identified morphologically as normal epithelial crypts. These cells are visualized using **outlining** in Fig 1b, where they form distinct circular structures in the tissue. The expert speculated that the elevated pan-cytokeratin signal was due to nonspecific antibody binding, noting that such details could not be inferred from the embedding and feature data alone. Fig 1 displays this set as visualized in all three views.

The expert noted the similarity between this set visualization and the heatmaps/dendrograms in the CyLinter paper [5], which hierarchically cluster cells (e.g., tumor, immune, stromal). They emphasized that supporting hierarchical grouping of sets would help further dissect the data. They also stressed the importance of quality control—manual inspection is often the only way to distinguish true findings from artifacts. Thus, enabling the omission of specific clusters or regions from analysis is essential.

### 7.2 Melanoma Analysis with a Cancer Biologist

Also in the field of oncology, we conducted two hour-long sessions with a cancer biologist in which we explored a tissue sample from a melanoma patient with **SEAL**. This expert has training in both computational biology and immuno-oncology and is interested in studying the spatial signatures of cancer, particularly in melanoma. The primary goal of their analysis was to identify distinct tumor morphologies that could serve as clinical biomarkers, focusing on mitotic cells, which indicate rapid tumor growth, and multinucleated cells, which are linked to poor prognosis. Traditional pathology relies on manual inspection of arbitrarily selected regions within whole slide images stained with hematoxylin and eosin (H&E), the common pink and purple histological stain that highlights cellular structures. While H&E staining remains the gold standard for tissue diagnosis, pathologists often work in highly variable and subjective ways. Their annotations and diagnostic decisions tend to reduce complex tissue morphology to binary outcomes, making it challenging to standardize analysis and develop computational methods that leverage the rich information contained within tissue architecture.

This specimen, a whole-slide section of melanoma tissue, was collected using an imaging technique that collects both H&E and high-plex immunofluorescence images on the same tissue, enabling the integration of single-cell analysis, as outlined in the prior case study (Sec 7.1), into the standard histopathological workflow. More specifically, the expert is investigating whether Variational Autoencoders (VAE), trained on pixel-level data from cells imaged using both H&E staining and multiplex immunofluorescence, can create a shared latent representation that bridges traditional histopathological practice and single-cell molecular analysis. Cells were encoded into a 128-dimensional latent space and then reduced to a two-dimensional embedding for visualization using UMAP, with hyperparameters iteratively tuned to enhance biological interpretability. The VAE reconstructed individual cells and created a structured embedding of their features, enabling exploration and clustering of this embedding as a means to uncover biologically and clinically relevant patterns. The biologist is particularly interested in dividing (mitotic) cells to examine if spatial patterns or single-cell features correlate with the proliferation of tumor cells. Additionally, the study investigated neutrophils, a type of white blood cell important for fighting infections. These cells lack molecular markers but may have diagnostic significance in oncology.

The biologist began by toggling the red, green, and blue channels of the H&E stain. This widely used technique enhances contrast between cell nuclei and surrounding tissue structures, providing clear insights into tissue architecture and pathology. In the **Hybrid Embedding** view (Fig 8), individual cells and their nuclei became evident. The biologist focused on a region in the embedding where individual cells appeared distinctly pinker than those elsewhere. Direct visual inspection of these cells in the **Hybrid Embedding** revealed clear morphological signs of necrosis, including intensely pink-stained cytoplasm and sharply faded or absent nuclei, distinguishing them from surrounding viable cells. This observation strongly highlights the power of combining morphological detail captured by H&E staining with learned molecular embeddings, enabling precise identification of critical tissue features and substantially enhancing biological interpretation and clinical insight. Next, the biologist examined the k-means clusters in the **Set View** and quickly noted that SOX10, a tumor marker, was consistently important across clusters (Fig 8), mirroring the findings of Case Study 1 (Sec 7.1). Moving to the **Hybrid Embedding** view, they explored the spatial expression patterns of immune markers, CD4, which is found primarily on Helper T cells, and CD8, which is found on Cytotoxic T cells. Cells expressing either CD4 or CD8 appeared in the same localized region of the embedding, suggesting the model grouped them based on shared immune-related features. These markers were not co-expressed within the same cells; rather, they were expressed in distinct but closely embedded cells, indicating a potential shared functional role or tissue context captured by the VAE. When this region of interest was highlighted in the **Spatial Image View**, the CD4- and CD8-expressing cells were found to be spatially adjacent at the tumor boundary, consistent with known patterns of immune infiltration. This suggested that the VAE was capturing some biologically meaningful properties, even though it did not differentiate strongly between the two T cell subtypes in the embedding space. Finally, the expert examined the presence of Phospho-H3 (a mitotic marker) and Ki67 (a proliferation marker) within the **Hybrid Embedding** to assess whether actively dividing cells were forming distinct clusters. However, no clear separation emerged, suggesting that the current model may not yet capture the subtle features associated with these biological states. Similar to the CD4 and CD8 cluster, this highlights the ability of **SEAL** to serve as a tool for model inspection, which in turn supports human-in-the-loop model development and refinement. The biologist in this case study has since been set up with **SEAL** to support ongoing exploration as they incrementally modify their data preprocessing steps, VAE model architecture, and training regimen, with the goal of identifying markers or morphological signatures that explain these more ambiguous cell populations. This process not only aims to reveal markers linked to aggressive tumor behavior, but also to deepen our understanding of how cancer spreads and responds to therapy.

**Figure 8.**
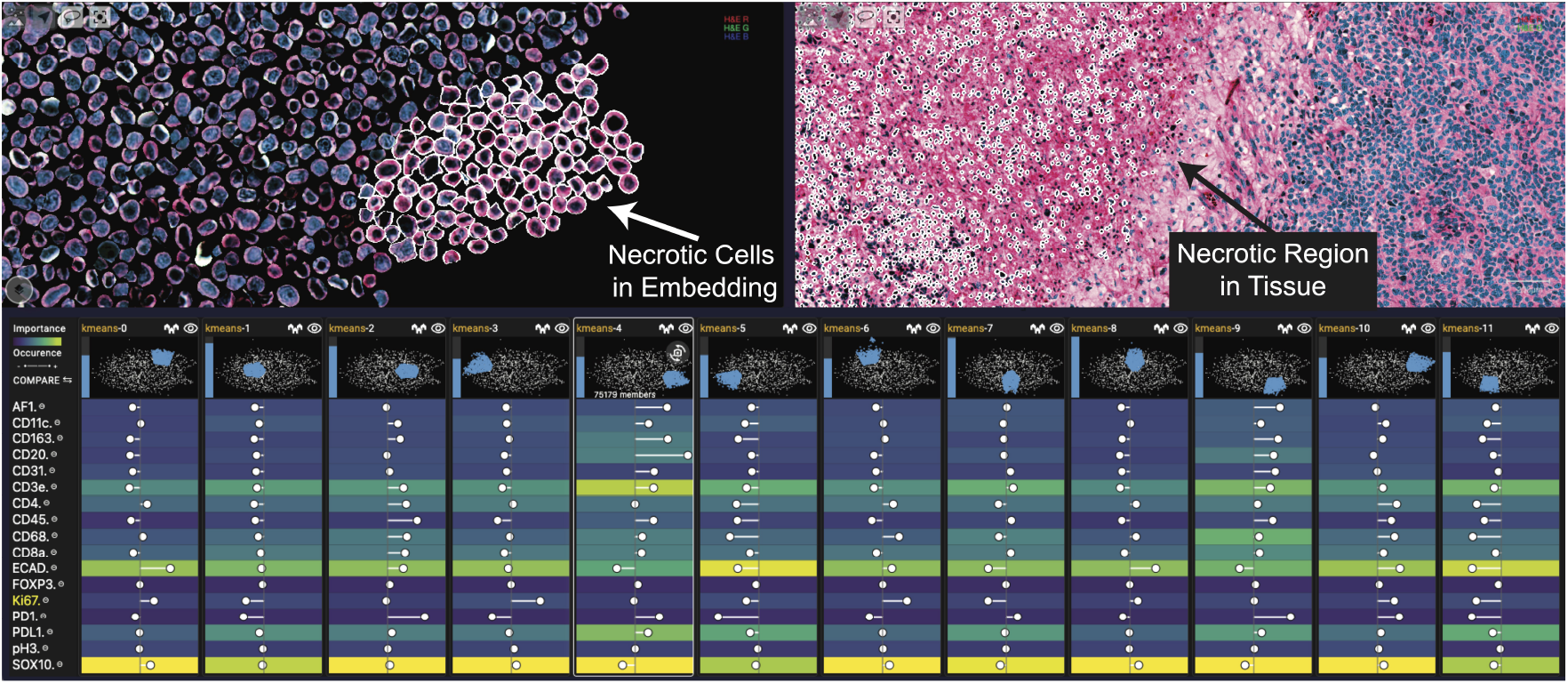
Case Study 2: Melanoma Analysis. The cell morphology of the selected cluster, when visualized with H&E, reveals visually discrete cells in the Hybrid Embedding. These cells are necrotic and correspond to a discrete necrotic region in the Spatial Image.

### 7.3 Astronomical Analysis

We conducted a separate case study on astronomical data with a postdoctoral fellow at the Harvard–Smithsonian Center for Astrophysics with over a decade of experience in developing and maintaining tools for the radio astronomy community. In the field of astronomy, the Sloan Digital Sky Survey (SDSS) [1] remains one of the most widely analyzed and well understood datasets due to its data documentation and accessibility. Similar to tissue samples, SDSS images are composed of multiple image filters, commonly referred to as *ugriz*. Each filter captures light from a narrow wavelength. Stacking filters onto one another then creates a combined image of a single patch of sky across a wide spectrum from near ultraviolet to near infrared. Within this image, celestial objects, such as stars and galaxies, can be identified and segmented, in a process that is remarkably similar to how single cells are cut out from whole tissue images. During our one-hour study session with the expert, we demonstrated how a single field of SDSS data is visualized in **SEAL**, and explored whether our tool could validate well-known patterns in astrophysics. From the **Set View** the expert immediately noticed that some known oddities show up clearly in the visualization, such as the importance of a given band (Fig 9, left). The *z*-filter, for instance, captures the band of light farthest into the near-infrared wavelength. As such, it tends to be the noisiest or contain the least distinct data. This shows up in the feature importance heatmap in **SEAL**, where the *z* filter was the least important in distinguishing any of the clusters.

**Figure 9.**
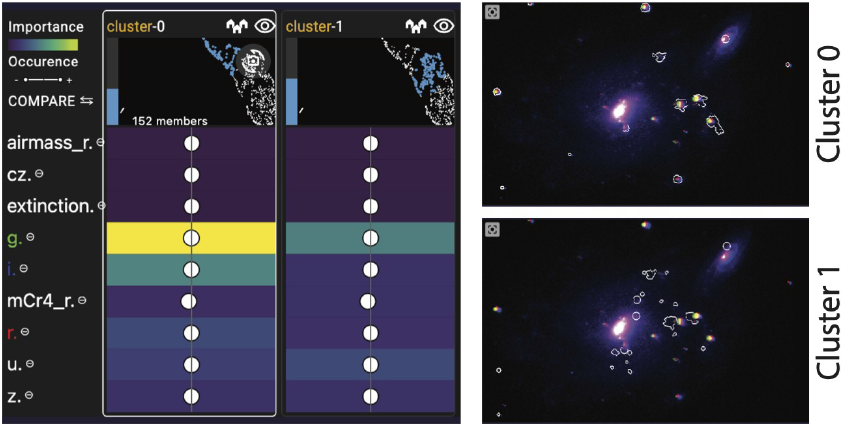
Case Study 3: Investigating astronomical imaging data from the Sloan Digital Sky Survey with an astrophysicist using **SEAL**. The **Spatial Image View** depicts two galaxies, NGC 450 and UGC 807.

Next, from the **Hybrid Embedding** view and the **Spatial Image View**, the expert was able to identify and explain the presence of two distinct types of stars in galaxies. The image depicted two spiral galaxies (Fig 9, right), NGC 450 and UGC 807. It is known that spiral galaxies are younger and still actively forming stars in their spiral arms. These young stars burn hotter than older stars and tend to emit blue light, resulting in the galaxy arms appearing more blue and less red. In contrast, the galaxy centers have less star formation. This means that they no longer have massive, young stars and tend to take on the red hue of older stars. In the **Spatial Image**, the expert could identify this expected pattern, with the bright galaxy centers surrounded by a blue halo. Interestingly, this image also shows large stars in the vicinity of the galaxy. These stars are not part of the galaxy, but only appear close because their depth has been compressed. This difference can be seen in the **Hybrid Embedding** view and the **Set View**, where Cluster 1 captures entities in the galaxy, while the large stars are grouped into a separate Cluster 0 (Fig 9). From the **Set View**, the expert further elaborated that because the *gri* filters are the deeper ones, their feature importance scores indicate two things: the age of stars in the galaxy, and also a continuous distribution of distances because the redshift effect causes light emitted by distant objects to be shifted towards a longer wavelength, and thus appear more red. They suggested that the importance of some of the filters, such as *g* and *i*, are a reflection of these two phenomena.

Overall, while the expert agreed that the SDSS data used in this case study effectively showcased many of the features of **SEAL** and revealed patterns that aligned well with established astrophysics knowledge, they also pointed out that data from a single patch of sky was too limited to fully utilize the **Spatial Image** view. However, they were excited about the potential for **SEAL** to reveal large-scale structures such as voids, clusters, and filaments that only show up in images at the most massive scales. Another potential application of the **Spatial Image** view would be to investigate galaxy interactions for nearby galaxies; when a galaxy falls into a main cluster, it starts to lose its gas, which gets stripped off by the strong radiation coming from the center region. This would likely look different from a galaxy that has come through the cluster once and is now on its orbit back out. As such, by combining the **Spatial Image** and **Set View**, the expert anticipated that the visualizations would reveal trends in multi-galaxy interactions that are otherwise more difficult to determine. Overall, the expert expressed enthusiasm for **SEAL**, and suggested future collaborations with more extensive datasets for exploration.

## 8 Discussion and Limitations

From our case studies, domain experts broadly found **SEAL** to be useful for characterizing clusters, investigating the spatial origins of unexpected patterns in the embedding space, explaining and refining models such as VAEs, and validating set features against known scientific facts. However, they also surfaced some limitations of the tool and opportunities for future work.

### 8.1 Enhancing the Embedding Space

We found that analysts in our case studies consistently prioritized the **Hybrid Embedding** space as their primary analytic lens, using the **Spatial Image** view primarily to contextualize and assign meaning to clusters within it. As such, we argue that future research should focus on strengthening this view rather than seeking to replace it. This entails addressing well-known limitations of DR techniques. In **SEAL**, we take a step in this direction by enriching the embedding space with spatial and image information and mitigating occlusion through the **Hybrid Embedding** view. However, we believe there are still unexplored opportunities for improving embedding spaces. Algorithms like UMAP and t-SNE focus largely on preserving local structure, and while UMAP [54] aims to retain global structure, empirical support remains mixed [33, 85]. This ambiguity highlights the need for visualization tools that make such limitations transparent to analysts. Similarly, Case Study 2 (Sec 7.2) highlights the necessity of comparing results across algorithms or hyperparameter settings, particularly when users are interested in improving or refining the algorithm.

One persistent challenge is the stochastic nature of many DR methods, which can yield inconsistent embeddings from the same data [48, 54]. Additionally, as seen in Case Study 1 (Sec 7.1) and detailed in Baker et al. [5], sensitivity to artifacts and outliers further complicates analysis. Comparing embeddings and incorporating new data remains a challenging task due to these instabilities. To advance the utility of embedding-based analysis, we call for robust visualization methods that not only reveal the variability and fragility of DR algorithms but also offer principled approaches to overcome them.

### 8.2 Formalizing Relationships Between Sets

In Case Study 1 (Sec 7.1), we also examined how organizing sets into hierarchies provides a powerful framework for dissecting complex data. In this context, such hierarchies capture the stratification of cell types, beginning with broad categories—such as tumor, immune, or stromal—and progressing toward more specific subtypes and cellular states. The relationships between biological entities often extend beyond simple hierarchical structures, encompassing more intricate patterns such as spatial organization or overlapping functional roles. These richer, ontological relationships were a key motivation for developing **SEAL**. We therefore argue that **SEAL** should be extended to support not only hierarchical relationships but also more complex, network-like associations between sets. This entails an element of network analysis and visualization, as domain experts aim to formalize and explore these layered ontologies. In this broader view, the **Hybrid Embedding** and **Set View** can work in tandem to represent complex set relationships: for example, hierarchies might be visualized as smaller sets nested within larger ones. Nonetheless, such visualizations may not always align with domain-specific conventions—for instance, biomedical hierarchies are often represented explicitly as trees. As such, aligning visualization methods with established ontologies remains an important challenge in supporting expert interpretation.

### 8.3 Scaling up the Dimensions

The spatial imaging and tabular feature datasets analyzed in **SEAL** are broadly applicable across domains, as shown in our case studies. The tool assumes the existence of specific data formats: spatial information is encoded in a 2D image, tabular features are scalar, and a DR algorithm maps these scalars into a 2D embedding. The surrogate feature importance model also operates in this 2D space. While such simplifications aid tractability and computational efficiency, they may overlook the full physical complexity of the underlying systems.

Consider, for example, that spatial information is inherently volumetric, with objects occupying 3D space and possessing meaningful positions along the z-axis. The SDSS data in Case Study 3 (Sec 7.3), for instance, compresses the depth of celestial objects such that they appear on the same plane in the **Spatial Image**. However, the lack of depth can be misleading, making distant objects appear close together—the two galaxies in Fig 9 are in reality separated by a distance of more than 400 million lightyears. Likewise, tabular features may include vector quantities, such as the velocity of moving entities. Finally, spatial datasets may also include temporal dimensions, where sequential snapshots reflect how entities evolve over time. In astronomy, celestial bodies follow orbital paths and undergo dynamic events like supernovae; in biomedical contexts, immune cells become activated or depleted as they battle antigens. Thus, as data acquisition methods—such as 3D imaging, motion tracking, and time-resolved measurements—continue to advance across disciplines, it is increasingly important to extend **SEAL** to support visualizations and analyses that more faithfully reflect the multidimensional and dynamic nature of real-world systems.

## 9 Conclusion

In summary, most current analysis techniques for spatial datasets either emphasize high-level abstractions—such as the outputs of dimensionality reduction algorithms or overall tissue functions—or focus on low-level details like individual object morphologies. However, an enduring challenge remains: connecting these high-level abstractions with the nuanced, low-level spatial details inherent in the original data. For instance, biomedical researchers often observe intriguing mesoscale features in tissue imaging that can be described by low-level multidimensional data attributes, yet no existing methods accurately reconstruct the intricate form and function of the original tissue solely from these data attributes. This gap highlights the critical role of spatial information, which preserves essential details lost in conventional high-dimensional representations and their DR abstractions. **SEAL** addresses the need to bridge the two levels of analysis by integrating spatial information directly into embedding spaces to support more informed, human-driven investigations. Moving forward, visualization tools like **SEAL** have the potential to deepen our understanding of how low-level spatial details contribute to high-level emergent behaviors in complex spatial datasets, ultimately guiding the development of more effective automated analysis methods that lessen our reliance on manual exploration.

## 10 Acknowledgments

This work was partially supported by NIH grants U01-CA284207 and R01HD104969.

## Footnotes

1 https://github.com/labsyspharm/cellcutter

## References

[1] Abdurro’uf K. Accetta, et al. The Seventeenth Data Release of the Sloan Digital Sky Surveys: Complete Release of MaNGA, MaStar, and APOGEE-2 Data. The Astrophysical Journal Supplement Series, 259:35, Apr. 2022. 1, 2, 3, 8

[2] B. Alper, N. Riche, et al. Design Study of LineSets, a Novel Set Visualization Technique. IEEE Transactions on Visualization and Computer Graphics, 17(12):2259–2267, Dec. 2011. 3

[3] B. Alsallakh, W. Aigner, et al. Radial Sets: Interactive Visual Analysis of Large Overlapping Sets. IEEE Transactions on Visualization and Computer Graphics, 19(12):2496–2505, Dec. 2013. 3

[4] J. Babbar and N. Rathee. Satellite Image Analysis: A Review. In 2019 IEEE International Conference on Electrical, Computer and Communication Technologies (ICECCT), pages 1–6, Feb. 2019. 1, 3

[5] G. J. Baker, E. Novikov, et al. Quality control for single-cell analysis of high-plex tissue profiles using CyLinter. Nature Methods, 21(12):2248–2259, Dec. 2024. 7, 10

[6] E. Becht, L. McInnes, et al. Dimensionality reduction for visualizing single-cell data using UMAP. Nature Biotechnology, 37(1):38–44, Jan. 2019. 1

[7] A. Boggust, B. Carter, and A. Satyanarayan. Embedding comparator: Visualizing differences in global structure and local neighborhoods via small multiples. In Proceedings of the 27th international conference on intelligent user interfaces, pages 746–766, 2022. 2

[8] T. Cabezon Pedroso and D. Byrne. Browsing the Latent Space: A New Approach to Interactive Design Exploration for Volumetric Generative Systems. In Proceedings of the 15th Conference on Creativity and Cognition, C&amp;C ‘23, pages 330–333, New York, NY, USA, Jun. 2023. Association for Computing Machinery. 2

[9] A. Chatzimparmpas, R. M. Martins, et al. The State of the Art in Enhancing Trust in Machine Learning Models with the Use of Visualizations. Computer Graphics Forum, 39(3):713–756, 2020. 4

[10] H. Chung, S. Nandhakumar, and S. Yang. GridSet: Visualizing Individual Elements and Attributes for Analysis of Set-Typed Data. IEEE Transactions on Visualization and Computer Graphics, 28(8):2983–2998, Aug. 2022. 3, 6

[11] A. O. Clarke, A. M. M. Scaife, et al. Identifying galaxies, quasars, and stars with machine learning: A new catalogue of classifications for 111 million SDSS sources without spectra. Astronomy & Astrophysics, 639:A84, Jul. 2020. 1

[12] A. Coenen and A. Pearce. Understanding UMAP, 2019. 2

[13] C. Collins, G. Penn, and S. Carpendale. Bubble Sets: Revealing Set Relations with Isocontours over Existing Visualizations. IEEE Transactions on Visualization and Computer Graphics, 15(6):1009–1016, Nov. 2009. 3

[14] T. L. Cook, B. Bandi, et al. Wide Area VISTA Extra-galactic Survey (WAVES): unsupervised star-galaxy separation on the WAVES-Wide photometric input catalogue using UMAP and hdbscan. Monthly Notices of the Royal Astronomical Society, 535(3):2129–2148, Dec. 2024. 1

[15] S. Corbugy, T. Septon, et al. Insight-SNE: Understanding t-SNE Embeddings through Interactive Explanation. ESANN 2024 proceesdings, pages 303–308, 2024. 2, 3

[16] S. Coy, S. Wang, et al. Single cell spatial analysis reveals the topology of immunomodulatory purinergic signaling in glioblastoma. Nature Communications, 13(1):4814, Aug. 2022. 1, 3

[17] R. Cutura, C. Morariu, et al. Hagrid — Gridify Scatterplots with Hilbert and Gosper Curves. In Proceedings of the 14th International Symposium on Visual Information Communication and Interaction, VINCI ‘21, pages 1–8, New York, NY, USA, Nov. 2021. Association for Computing Machinery. 2, 3

[18] K. Dinkla, M. J. van Kreveld, et al. Kelp Diagrams: Point Set Membership Visualization. Computer Graphics Forum, 31(3pt1):875–884, 2012. 3

[19] G. Garcia-Zanabria, M. M. Raimundo, et al. CriPAV: Street-Level Crime Patterns Analysis and Visualization. IEEE transactions on visualization and computer graphics, 28(12):4000–4015, Dec. 2022. 1, 3

[20] R. O. Green, M. L. Eastwood, et al. Imaging spectroscopy and the airborne visible/infrared imaging spectrometer (AVIRIS). Remote sensing of environment, 65(3):227–248, 1998. 3

[21] F. Heimerl, C. Kralj, et al. embComp: Visual Interactive Comparison of Vector Embeddings. IEEE Transactions on Visualization and Computer Graphics, 28(8):2953–2969, Aug. 2022. 2

[22] A. Holzinger, A. Saranti, et al. Explainable AI methods-a brief overview. In International workshop on extending explainable AI beyond deep models and classifiers, pages 13–38. Springer, 2020. 6

[23] H. Jeon, M. Aupetit, et al. Distortion-Aware Brushing for Interactive Cluster Analysis in Multidimensional Projections, Jan. 2022. 2, 3

[24] H. Jeon, A. Cho, et al. ZADU: A Python Library for Evaluating the Reliability of Dimensionality Reduction Embeddings. In 2023 IEEE Visualization and Visual Analytics (VIS), pages 196–200, Oct. 2023. 2

[25] H. Jeon, H.-K. Ko, et al. Measuring and Explaining the Inter-Cluster Reliability of Multidimensional Projections. IEEE Transactions on Visualization and Computer Graphics, 28(1):551–561, Jan. 2022. 2

[26] H. Jeon, H.-K. Ko, et al. Uniform Manifold Approximation with Twophase Optimization. In 2022 IEEE Visualization and Visual Analytics (VIS), pages 80–84, Oct. 2022. 2

[27] H. Jeon, Y.-H. Kuo, et al. Classes are Not Clusters: Improving Label-Based Evaluation of Dimensionality Reduction. IEEE Transactions on Visualization and Computer Graphics, 30(1):781–791, Jan. 2024. 2

[28] H. Jeon, G. J. Quadri, et al. CLAMS: A Cluster Ambiguity Measure for Estimating Perceptual Variability in Visual Clustering. IEEE Transactions on Visualization and Computer Graphics, 30(1):770–780, Jan. 2024. 2, 3

[29] J. Jessup, R. Krueger, et al. Scope2screen: Focus+ context techniques for pathology tumor assessment in multivariate image data. IEEE transactions on visualization and computer graphics, 28(1):259–269, 2021. 4

[30] S. Jin, A. Endert, and C. Andris. SNoMaN: a visual analytic tool for spatial social network mapping and analysis. Cartography and Geographic Information Science, 0(0):1–19, Oct. 2023. 1, 3

[31] M. S. Keller, I. Gold, et al. Vitessce: integrative visualization of multimodal and spatially resolved single-cell data. Nature Methods, pages 1–5, Sept. 2024. 7

[32] J. Kim, S. Rustam, et al. Unsupervised discovery of tissue architecture in multiplexed imaging. Nature Methods, 19(12):1653–1661, Dec. 2022. 1

[33] D. Kobak and G. C. Linderman. Initialization is critical for preserving global data structure in both t-SNE and UMAP. Nature Biotechnology, 39(2):156–157, Feb. 2021. 2, 9

[34] R. Kosara, F. Bendix, and H. Hauser. Parallel Sets: interactive exploration and visual analysis of categorical data. IEEE Transactions on Visualization and Computer Graphics, 12(4):558–568, Jul. 2006. 3

[35] R. Krueger, J. Beyer, et al. Facetto: Combining unsupervised and supervised learning for hierarchical phenotype analysis in multi-channel image data. IEEE transactions on visualization and computer graphics, 26(1):227–237, 2019. 3, 4

[36] L. Giovannangeli, F. Lalanne, et al. Guaranteed Visibility in Scatterplots with Tolerance. IEEE Transactions on Visualization and Computer Graphics, 30(1):792–802, Jan. 2024. 2, 3

[37] F. Lan, M. Young, et al. Visualization in Astrophysics: Developing New Methods, Discovering Our Universe, and Educating the Earth. Computer Graphics Forum, 40(3):635–663, 2021. 4

[38] A. Lex, N. Gehlenborg, et al. UpSet: Visualization of Intersecting Sets. IEEE Transactions on Visualization and Computer Graphics, 20(12):1983–1992, Dec. 2014. 3, 6

[39] M. Li, F. Choudhury, et al. ConcaveCubes: Supporting Cluster-based Geographical Visualization in Large Data Scale. Computer Graphics Forum, 37(3):217–228, 2018. 3

[40] Q. Li, K. S. Njotoprawiro, et al. EmbeddingVis: A Visual Analytics Approach to Comparative Network Embedding Inspection. In 2018 IEEE Conference on Visual Analytics Science and Technology (VAST), pages 48–59, Oct. 2018. 2

[41] Z. Li, T. Wang, et al. Construct boundaries and place labels for multiclass scatterplots. Journal of Visualization, 25(2):407–426, Apr. 2022. 2, 3

[42] J.-R. Lin, B. Izar, et al. Highly multiplexed immunofluorescence imaging of human tissues and tumors using t-CyCIF and conventional optical microscopes. eLife, 7:e31657, Jul. 2018. 1, 3

[43] J.-R. Lin, S. Wang, et al. Multiplexed 3D atlas of state transitions and immune interaction in colorectal cancer. Cell, 186(2):363–381, 2023. 7

[44] Y. Liu, E. Jun, et al. Latent Space Cartography: Visual Analysis of Vector Space Embeddings. Computer Graphics Forum, 38(3):67–78, 2019. 2

[45] S. M. Lundberg, G. G. Erion, and S.-I. Lee. Consistent individualized feature attribution for tree ensembles. arXiv preprint 1802.03888, 2018. 6

[46] S. M. Lundberg and S.-I. Lee. A unified approach to interpreting model predictions. In I. Guyon, U. V. Luxburg, et al., editors, Advances in neural information processing systems, volume 30. Curran Associates, Inc., 2017. 6

[47] X. Luo, L. Scandolo, and E. Eisemann. Texture Browser: Feature-based Texture Exploration. Computer Graphics Forum, 40(3):99–109, 2021. 2

[48] L. v. d. Maaten and G. Hinton. Visualizing Data using t-SNE. Journal of Machine Learning Research, 9(86):2579–2605, 2008. 1, 2, 10

[49] T. Manz, N. Abdennur, and N. Gehlenborg. anywidget: reusable widgets for interactive analysis and visualization in computational notebooks. Journal of Open Source Software, 9(102):6939, 2024. 7

[50] T. Manz, F. Lekschas, et al. A General Framework for Comparing Embedding Visualizations Across Class-Label Hierarchies. IEEE Transactions on Visualization and Computer Graphics, pages 1–11, 2024. 2, 3

[51] W.E. Marcílio-Jr, D. M. Eler, et al. HUMAP: Hierarchical Uniform Manifold Approximation and Projection. IEEE Transactions on Visualization and Computer Graphics, pages 1–10, 2024. 2

[52] Martín Abadi, Ashish Agarwal, et al. TensorFlow: Large-scale machine learning on heterogeneous systems, 2015. 2

[53] A. Mayorga and M. Gleicher. Splatterplots: Overcoming Overdraw in Scatter Plots. IEEE Transactions on Visualization and Computer Graphics, 19(9):1526–1538, Sept. 2013. 2

[54] L. McInnes, J. Healy, and J. Melville. Umap: Uniform manifold approximation and projection for dimension reduction. arXiv preprint 1802.03426, 2018. 1, 2, 9, 10

[55] F. Miranda, M. Hosseini, et al. Urban Mosaic: Visual Exploration of Streetscapes Using Large-Scale Image Data. In Proceedings of the 2020 CHI Conference on Human Factors in Computing Systems, CHI ‘20, pages 1–15, New York, NY, USA, Apr. 2020. Association for Computing Machinery. 1, 3

[56] C. Molnar. Interpretable machine learning. A guide for making black box models explainable. 3 edition, 2025. 6

[57] B. Montambault, G. Appleby, et al. DimBridge: Interactive Explanation of Visual Patterns in Dimensionality Reductions with Predicate Logic. IEEE Transactions on Visualization and Computer Graphics, pages 1–11, 2024. 2, 3

[58] M. Moor, M. Horn, et al. Topological Autoencoders. In Proceedings of the 37th International Conference on Machine Learning, pages 7045–7054. PMLR, Nov. 2020. 2

[59] J. Moore, C. Allan, et al. OME-NGFF: a next-generation file format for expanding bioimaging data-access strategies. Nature Methods, 18(12):1496–1498, Dec. 2021. 7

[60] R. Motta, R. Minghim, et al. Graph-based measures to assist user assessment of multidimensional projections. Neurocomputing, 150:583–598, Feb. 2015. 2

[61] A. J. Nirmal, Z. Maliga, et al. The Spatial Landscape of Progression and Immunoediting in Primary Melanoma at Single-Cell Resolution. Cancer Discovery, 12(6):1518–1541, Jun. 2022. 1, 3

[62] J. H. Park, S. Nadeem, and A. Kaufman. GeoBrick: exploration of spatiotemporal data. The Visual Computer, 35(2):191–204, Feb. 2019. 3

[63] K. Pearson. LIII. On lines and planes of closest fit to systems of points in space. The London, Edinburgh, and Dublin philosophical magazine and journal of science, 2(11):559–572, 1901. 1, 2

[64] N. Pezzotti, T. Höllt, et al. Hierarchical Stochastic Neighbor Embedding. Computer Graphics Forum, 35(3):21–30, 2016. 2

[65] G. Polder and G. W. van der Heijden. Visualization of spectral images. In Y. Censor and M. Ding, editors, Visualization and optimization techniques, volume 4553, pages 132 – 137. SPIE / International Society for Optics and Photonics, 2001. 2

[66] S. K. N. Portillo, J. K. Parejko, et al. Dimensionality Reduction of SDSS Spectra with Variational Autoencoders. The Astronomical Journal, 160(1):45, Jun. 2020. 3

[67] G. J. Quadri, J. A. Nieves, et al. Automatic Scatterplot Design Optimization for Clustering Identification. IEEE Transactions on Visualization and Computer Graphics, 29(10):4312–4327, Oct. 2023. 2, 3

[68] H. Rave, V. Molchanov, and L. Linsen. De-cluttering Scatterplots with Integral Images. IEEE Transactions on Visualization and Computer Graphics, pages 1–13, 2024. 2, 3

[69] M. T. Ribeiro, S. Singh, and C. Guestrin. “Why should i trust you?” Explaining the predictions of any classifier. In Proceedings of the 22nd ACM SIGKDD international conference on knowledge discovery and data mining, pages 1135–1144, 2016. 6

[70] R. Sadana, T. Major, et al. OnSet: A Visualization Technique for Large-scale Binary Set Data. IEEE Transactions on Visualization and Computer Graphics, 20(12):1993–2002, Dec. 2014. 3, 6

[71] T. Sainburg, L. McInnes, and T. Q. Gentner. Parametric UMAP embeddings for representation and semisupervised learning. Neural Computation, 33(11):2881–2907, 2021. 6

[72] A. Sarikaya and M. Gleicher. Scatterplots: Tasks, Data, and Designs. IEEE Transactions on Visualization and Computer Graphics, 24(1):402–412, Jan. 2018. 5

[73] D. Schapiro, H. W. Jackson, et al. histoCAT: analysis of cell phenotypes and interactions in multiplex image cytometry data. Nature methods, 14(9):873–876, Sept. 2017. 3

[74] P. Simonetto, D. Auber, and D. Archambault. Fully Automatic Visualisation of Overlapping Sets. Computer Graphics Forum, 28(3):967–974, 2009. 3

[75] N. Smith and N. van der Walt. A Better Default Colormap for Matplotlib, 2015. 6

[76] J.-T. Sohns, M. Schmitt, et al. Attribute-based explanation of nonlinear embeddings of high-dimensional data. IEEE Transactions on Visualization and Computer Graphics, 28(1):540–550, 2021. 2

[77] A. Somarakis, V. Van Unen, et al. ImaCytE: Visual Exploration of Cellular Microenvironments for Imaging Mass Cytometry Data. IEEE Transactions on Visualization and Computer Graphics, pages 1–1, 2019. 3

[78] P. Strobl, A. A. Mueller, et al. Laboratory calibration and inflight validation of the digital airborne imaging spectrometer DAIS 7915. In Algorithms for multispectral and hyperspectral imagery III, volume 3071, pages 225–236. SPIE, 1997. 3

[79] J. B. Tenenbaum, V. d. Silva, and J. C. Langford. A Global Geometric Framework for Nonlinear Dimensionality Reduction. Science, 290(5500):2319–2323, Dec. 2000. 2

[80] J. Venna and S. Kaski. Neighborhood preservation in nonlinear projection methods: An experimental study. In Proceedings of the international conference on artificial neural networks, Icann ‘01, pages 485–491, Berlin, Heidelberg. 2001. Springer-Verlag. 6

[81] J. Venna and S. Kaski. Local multidimensional scaling. Neural Networks, 19(6):889–899, Jul. 2006. 2

[82] A. Vieth, B. Lelieveldt, et al. Interactions for Seamlessly Coupled Exploration of High-Dimensional Images and Hierarchical Embeddings. The Eurographics Association, 2023. 2

[83] A. Vieth, A. Vilanova, et al. Incorporating Texture Information into Dimensionality Reduction for High-Dimensional Images. In 2022 IEEE 15th Pacific Visualization Symposium (PacificVis), pages 11–20, Apr. 2022. 2

[84] J. Vihrovs, K. Prusis, et al. An Inverse Distance-based Potential Field Function for Overlapping Point Set Visualization:. In Proceedings of the 5th International Conference on Information Visualization Theory and Applications, pages 29–38, Lisbon, Portugal, 2014. SCITEPRESS - Science and and Technology Publications. 3

[85] S. Wang, E. D. Sontag, and D. A. Lauffenburger. What cannot be seen correctly in 2D visualizations of single-cell ‘omics data? Cell Systems, 14(9):723–731, Sept. 2023. 3, 9

[86] Z. J. Wang, F. Hohman, and D. H. Chau. WizMap: Scalable Interactive Visualization for Exploring Large Machine Learning Embeddings. arXiv 2306.09328, 2023. 2, 3

[87] S. Warchol, R. Krueger, et al. Visinity: Visual spatial neighborhood analysis for multiplexed tissue imaging data. IEEE transactions on visualization and computer graphics, 29(1):106–116, 2022. 3, 4, 6

[88] S. Warchol, J. Troidl, et al. psudo: Exploring multi-channel biomedical image data with spatially and perceptually optimized pseudocoloring. In Computer graphics forum, volume 43, page e15103. Wiley Online Library, 2024. 4

[89] C. Yapp, E. Novikov, et al. UnMICST: Deep learning with real augmentation for robust segmentation of highly multiplexed images of human tissues. Communications Biology, 5(1):1–13, Nov. 2022. 7

[90] J. Zhang, A. Malik, et al. Topogroups: Context-preserving visual illustration of multi-scale spatial aggregates. In Proceedings of the 2017 CHI conference on human factors in computing systems, pages 2940–2951, 2017. 3

[91] Y. Zhang, W. Luo, et al. Visualizing the Impact of Geographical Variations on Multivariate Clustering. Computer Graphics Forum, 35(3):101–110, 2016. 1, 3

